# A global analysis of song frequency in passerines provides no support for the acoustic adaptation hypothesis but suggests a role for sexual selection

**DOI:** 10.1101/2020.06.30.179812

**Authors:** Peter Mikula, Mihai Valcu, Henrik Brumm, Martin Bulla, Wolfgang Forstmeier, Tereza Petrusková, Bart Kempenaers, Tomáš Albrecht

**Affiliations:** Department of Zoology, Faculty of Science, Charles University, Viničná 7, 128 44, Praha 2, Czech Republic; Institute of Vertebrate Biology, Czech Academy of Sciences, Květná 8, 603 65, Brno, Czech Republic; Department of Behavioural Ecology and Evolutionary Genetics, Max Planck Institute for Ornithology, Eberhard-Gwinner-Str. 7, 82319 Seewiesen, Germany; Communication and Social Behaviour Group, Max Planck Institute for Ornithology, Eberhard-Gwinner-Str. 11, 82319 Seewiesen, Germany; Department of Ecology, Faculty of Environmental Sciences, Czech University of Life Sciences, 16521 Prague, Czech Republic; Department of Ecology, Faculty of Science, Charles University, Viničná 7, 128 43, Praha 2, Czech Republic

**Keywords:** acoustic adaptation hypothesis, allometry, animal communication, bird song, macroecology, morphological constraints, sexual selection

## Abstract

Many animals use acoustic signals for communication, implying that the properties of these signals can be under strong selection. The acoustic adaptation hypothesis predicts that species living in dense habitats emit lower-frequency sounds than those in open areas, because low-frequency sounds generally propagate further in denser vegetation. Signal frequency may also be under sexual selection, because it correlates with body size and lower-frequency sounds are perceived as more intimidating. Here, we evaluate these hypotheses by analysing variation in peak song frequency across 5,085 passerine species (Passeriformes). A phylogenetically-informed analysis revealed that song frequency decreases with increasing body mass and with male-biased sexual size dimorphism. However, we found no support for the predicted relationship between frequency and habitat. Our results suggest that the global variation in passerine song frequency is mostly driven by natural and sexual selection causing evolutionary shifts in body size rather than by habitat-related selection on sound propagation.

**Statement of authorship:** TA and PM conceived and designed the study with input from all authors. TA and BK coordinated the study. PM collected the song data. MV performed the statistical analyses with input from WF. MB made the figures with help from MV and PM. TP and HB provided bioacoustic expertise. PM drafted the first version of the manuscript. TA, BK and PM revised and finalized the manuscript with input from all authors.

**Data availability statement:** The data used in this study were collected from publicly available databases. All data and computer code used to generate the results, as well as supplementary figures and tables will be freely available at https://osf.io/fa9ky/.

## INTRODUCTION

Acoustic signalling is widespread among animals (Bradbury & Vehrencamp 1998; Gerhardt & Huber 2002; Catchpole & Slater 2008). Successful transmission and reception of acoustic signals between conspecifics are essential in diverse contexts, including predation avoidance (alerting others to a threat), territory defence, mate attraction, and synchronization of breeding activities (Bradbury & Vehrencamp 1998; Catchpole & Slater 2008). One of the fundamental characteristics of acoustic signals is the frequency of the sound, because it strongly affects signal propagation through the environment (Morton 1975; Wiley & Richards 1982; Padgham 2004). Low frequency sounds are generally less attenuated during transmission than high frequency sounds (Wiley & Richards 1982; Padgham 2004). Nevertheless, the frequency of acoustic signals is tremendously diverse across the animal kingdom (Gerhardt 1994; Fitch 2006; Gillooly & Ophir 2010; Pijanowski *et al.* 2011) and several hypotheses have been proposed to explain this diversity. Here, we focus on the three most compelling ones: (1) the acoustic adaptation hypothesis, (2) the morphological constraint hypothesis, and (3) the sexual selection hypothesis.

Since the 1970s, it has been postulated that the frequency of acoustic signals could reflect an adaptation to maximize the effectiveness of sound transmission in specific habitats (Morton 1975). This is known as the acoustic adaptation hypothesis (Boncoraglio & Saino 2007; Ey & Fischer 2009). Sounds transmitted through the natural environment are subject to degradation, for example due to environmental absorption, reverberation and scattering. The degree of this degradation depends both on the sound structure and on the physical characteristics of the environment (Wiley & Richards 1982; Brumm & Naguib 2009). Specifically, because of frequency-dependent attenuation, low-frequency sounds transmit generally further than high-frequency sounds. However, the slope of the frequency dependence is steeper in dense, forested habitats because of the high degree of sound absorption and scattering from foliage. Hence, high-frequency signals are attenuated more strongly in closed than in open habitats (Morton 1975; Marten & Marler 1977; Wiley & Richards 1978). Therefore, species living in forested habitats are expected to produce vocalizations of lower frequencies than those living in open habitats (Ey & Fischer 2009). Despite this strong theoretical underpinning, empirical evidence for the acoustic adaptation hypothesis is equivocal (Morton 1975; Wiley 1991; Buskirk 1997; Bertelli & Tubaro 2002; Blumstein & Turner 2005; Ey & Fischer 2009). For instance, a meta-analysis by Boncoraglio & Saino (2007) showed that song frequency in birds tends to be lower in closed compared with open habitats, but the effect size was small. A review by Ey & Fischer (2009) concluded that habitat-related adjustments of frequency parameters of acoustic signals of birds, anurans and mammals are not as widespread as previously thought.

The morphological constraint hypothesis simply posits that body size sets a limit on the frequency of the sound an animal can produce. Morphological constraints generally seem to play a pervasive role in the evolution of animal acoustic communication (Ryan & Brenowitz 1985; Bradbury & Vehrencamp 1998; Fitch & Hauser 2002). A negative relationship between body size and frequency of acoustic signals, i.e. larger species tend to produce lower frequency sounds than smaller species, seems to be a general rule in animal bioacoustics and has been documented across various groups, including insects, fishes, amphibians, reptiles, birds, and mammals (Wallschläger 1980; McClatchie *et al.* 1996; Fitch & Hauser 2002; Gillooly & Ophir 2010; Pearse *et al.* 2018). In birds, it has been suggested that the frequency of vocalizations negatively scales with body size, simply because body size influences the morphology and functional aspects of the vocal apparatus, such as the size of vibratory structures (Bertelli & Tubaro 2002; Suthers & Zollinger 2008; Seneviratne *et al.* 2012; Gonzalez-Voyer *et al.* 2013; Tietze *et al.* 2015). However, body size alone does not explain the entire variation in song frequency across animals. Departures from the negative allometric relationship between frequency of acoustic signals and body size may reflect (a) differences in evolutionary history that caused variation in syrinx or vocal tract morphology (phylogenetic constraints) and (b) differences in costs or benefits of producing low-frequency sounds. Thus, variation in frequency may inform about current or past selection on acoustic signals (Searcy & Nowicki 2005; Ophir *et al.* 2010; Wagner *et al.* 2012).

This brings us to the hypothesis that the frequency of acoustic signals may be sexually selected, acting as an indicator of an individual’s size, dominance or fighting ability. In various taxa, the frequency of male vocalizations indeed seems to indicate individual body size and can influence territory establishment (or other forms of male-male competition), attractiveness (female choice) and ultimately an individual’s reproductive success (Morton 1977; Fitch & Hauser 2002; Apicella *et al.* 2007; Hardouin *et al.* 2007; Mager *et al.* 2007; Vannoni & McElligott 2008; Forstmeier *et al.* 2009; Brumm & Goymann 2017). For instance, the frequency of advertising vocalizations negatively correlates with body size in males of common toads *Bufo bufo* and during the mating period smaller males were less often attacked by larger males when natural croaks of the small males were experimentally replaced by deep croaks (Davies & Halliday 1978). Similarly, heavier individuals of scops owl *Otus scops* produced lower-frequency hoots and territorial males responded less intensely to hoots simulating heavier intruders (Hardouin *et al.* 2007). Thus, if low-frequency sounds are advantageous during agonistic interactions between males and as a means of dominance status signalling (Davies & Halliday 1978; Wagner 1989; Briefer *et al.* 2010; Bro-Jørgensen & Beeston 2015), we predict correlated evolution of male vocal frequency and indices of the intensity of sexual selection such as male-biased sexual size dimorphism (Trivers 1972; Fairbairn 1997).

Here, we use a large data set of 5,085 passerine species (Order: Passeriformes), representing 85% of all passerines and 50% of all avian taxa (Jetz *et al.* 2012), to explore interspecific variation in peak frequency of male song. Applying a phylogenetically-informed cross-species analysis, we evaluate the association between song frequency and habitat density, body size (expressed as body mass), and the intensity of sexual selection (expressed as sexual size dimorphism). Based on the hypotheses outlined above, we test the one-tailed predictions that lower-frequency songs are associated with (1) more closed (forested) habitats, (2) larger body size and (3) stronger male-biased sexual size dimorphism.

Passerines are an excellent study system for evaluating sources of interspecific variation in signal frequency. First, their song represents a textbook example of a long-range acoustic signal that plays an important role in mate attraction and territory defence (Catchpole 1987; Catchpole & Slater 2008). Second, passerines are globally distributed, show a more than 300-fold difference in body mass, vary in sexual selection pressures and mating systems, and occupy a wide range of habitats (del Hoyo *et al.* 2018). Although song (or call) frequency has been widely studied in birds, previous comparative studies often evaluated the effects of body size, sexual selection, and habitat effects separately and without accounting for phylogeny (reviewed by Ey & Fischer 2009). Moreover, previous studies were restricted to a few species only (Ey & Fischer 2009).

## MATERIALS AND METHODS

### Data on peak song frequency

We collected song recordings primarily from xeno-canto (https://www.xeno-canto.org), a citizen science repository of bird vocalizations. When access to recordings of endangered or vulnerable species was restricted, we directly contacted the authors. For species with missing recordings on xeno-canto, we used recordings from the Macaulay Library (The Cornell Lab of Ornithology, https://www.macaulaylibrary.org/). We focused exclusively on the song, ignoring other types of vocalizations (e.g. calls). Song is commonly defined as a long-range vocalization that is used mainly in mate attraction and territory defence. The definition of the song may, however, vary across sources or authorities, and functions of particular vocalizations are still poorly known for several passerine species. Therefore, we used the classification of vocalizations as provided on the platform storing the recordings. Although some recordings might be misclassified, we primarily focused on high-quality recordings (scored as quality “A” or “B” in xeno-canto, or rated four or more stars in Macaulay Library), usually collected by skilled observers with in-depth knowledge of particular bird species’ vocalizations. Both repositories also provide a space for discussion and correction of misclassified recordings by community members, increasing the reliability of the available information.

We collected 1-5 (median = 4, mean ± SD = 3.7 ± 1.5) recordings of adult male song for each species (total of 18,789 recordings from 5,085 species). We did not use recordings of female and juvenile song. However, recordings often lacked information on sex, age, or the number of singing individuals. Although most of such recordings presumably documented adult male song, females of many species sing, either solo, in duets (coordinated joint singing of a mated pair) or in a chorus (three and more singing individuals) (Odom *et al.* 2014; Tobias *et al.* 2016; Mikula *et al.* 2020). A few recording annotations mentioned “duet” or “chorus” and in some cases we could disentangle parts produced by different individuals. We then measured song frequency for the individual producing the more complex song, i.e. containing more elements and syllable types (presumably a male). For a few species, we were not able to separate the song of multiple individuals. In these cases, we assumed that the recording was representative of the song of the males of the species. Although this procedure might have introduced some error, we do not expect systematic bias in species-specific frequency values. We assigned geographic coordinates to all song recordings as reported by the person who made the recording. In widely distributed species, recordings were typically separated by tens to thousands of kilometres. However, in species with smaller ranges, we used recordings made at least 1 km apart to reduce the possibility that two or more analysed recordings contained song of the same individual. In several species (all island or mountain endemics or poorly sampled species) this was not possible. In these cases, we a priori maximized the altitudinal and temporal separation of recordings, by only selecting recordings that differed in altitude by at least 100 metres or were collected in different years.

After downloading, all recordings were converted to *.wav* format with an online converter (www.online-audio-converter.com) at a sampling rate of 44.1 kHz. We characterized song frequency by a single parameter, namely peak frequency (i.e. the frequency at maximum amplitude), using the Raven Pro 1.4 software (Cornell Lab of Ornithology, Ithaca, NY, USA, www.ravensoundsoftware.com). We then calculated the median value for each species. Peak frequency is central to our hypotheses because: (1) unlike minimum and maximum frequencies, it is crucial for signal transmission (Brumm & Naguib 2009), (2) it may differ between habitats (see meta-analysis in Boncoraglio & Saino 2007), and (3) it is a key trait in other studies investigating the effect of morphological constraints and sexual selection on acoustic communication (Gillooly & Ophir 2010; Greig *et al.* 2013; Mason & Burns 2015; Thiagavel *et al.* 2017). First, we measured peak song frequency based on a fast Fourier transform length of 256 points (Hann window), resulting in a frequency resolution of 172 Hz. In a second step, we re-measured peak song frequency for species with median peak frequency < 1.2 kHz *(n* = 90 species), using a higher frequency resolution of 21.5 Hz (fast Fourier transform length of 2,048 points) to capture the lower end of the range in peak song frequency more accurately. To ensure consistency, all recordings were downloaded and analysed by a single person (PM).

### Predictor variables

#### Body size and sexual size dimorphism

As a proxy of species-specific body size, we used mean body mass (in grams; pooling sexed and unsexed individuals from Dunning 2008; *n* = 4,602 species) or male body mass (from Dunning 2008; *n* = 984 species). To estimate sexual size dimorphism we used data on male and female body mass (from Dunning 2008; *n* = 984 species) or wing length (in millimetres; from Dale *et al.* 2007; *n* = 2,463 species). We then calculated sexual size dimorphism either as log(male body mass) – log(female body mass) or as log(male wing length) – log(female wing length). Positive values indicate species where males are larger than females, i.e. male-biased sexual size dimorphism. Sexual size dimorphism is associated with other indices of the intensity of sexual selection, such as the mating system (polygyny versus monogamy) or testis size (Dunn *et al.* 2001).

#### Habitat density

As a proxy for habitat density, we used tree cover data from Collection 2 of the Copernicus Global Land Cover project (Buchhorn *et al.* 2020). For each geographic location of a song recording, we extracted the percentage of tree cover in a 100 × 100 metres quadrant using the *exactextractr* package (v.0.2.1) in R (Baston 2020). Species-specific tree cover was then estimated as the mean of all conspecific recordings.

We also extracted data on habitat type for each species based on descriptions in del Hoyo *et al.* (2018). We assigned each species to the most prevalent habitat type on a three-point scale: (1) closed (covering species living in densely vegetated habitat types such as forest, woodland and mangrove), (2) mixed (covering generalist species and species inhabiting ecotones), and (3) open (covering species inhabiting grassland, steppe, desert and semi-desert, savannah, bushland, rocky habitats and seashores).

### Statistical analyses

All statistical analyses were performed using R v. 4.0.0 (R Development Core Team 2019).

#### Data visualization

To help interpret the investigated relationships, we assessed whether peak song frequency evolved within diverged groups of passerines by plotting the evolutionary tree of song frequency, as well as of the predictors (Fig. S1). We mapped these variables on a maximum credibility tree reconstructed from 100 trees using the function maxCladeCred in the *phangorn* package (v. 2.5.5) (Schliep 2011). Character states at internal nodes were mapped using a maximum-likelihood approach implemented in the contMap function (Revell 2013) from the *phytools* package (Revell 2012). To illustrate the geographic distribution of peak song frequency, we used the breeding range distribution of all passerines (obtained from BirdLife International and NatureServe 2018) to visualize mean peak song frequency values across passerine assemblages with grid cells of 112.5 × 112.5 km (~1° scale) (Valcu *et al.* 2012).

#### General modelling procedures

All comparative analyses were performed using the *phylolm* package (v. 2.6) (Tung Ho & Ané 2014). To control for non-independence due to common ancestry (Paradis 2011), we used phylogenetic generalized least-squares (PGLS) regressions with Pagel’s lambda (λ) transformation of a correlation structure (Pagel 1999). This method explicitly models how the covariance between species declines as they become more distantly related. If λ = 1, modelled traits co-vary in direct proportion to shared evolutionary history, whereas λ = 0 indicates phylogenetic independence of traits (Freckleton *et al.* 2002). We randomly sampled 100 phylogenetic trees (Hackett backbone) from those available at http://birdtree.org (Jetz *et al.* 2012), which included all species in our data set. We ran all models using these 100 phylogenies to account for uncertainties associated with different tree topologies and combined model coefficients by model averaging (Symonds & Moussalli 2011). For each model, we also calculated the proportion of variance explained (R^2^) according to Ives (2019) using the *rr2* package (Ives & Li 2018), including the conditional R2 (the variance explained by fixed and random effects) and the marginal R2 (the variance explained by the fixed effects only), and report these as mean values from 100 models each based on a different phylogenetic tree. Model residuals revealed no major violation of the assumptions of normality and homogeneity of variance. Peak song frequency and body mass were log-transformed before analysis. Peak song frequency and all predictors were also mean-centred and divided by their standard deviation (Schielzeth 2010).

Sex-specific body mass and wing length data were only available for 984 and 2,463 species, respectively. Hence, we estimated the missing values with the phylogenetic imputation method in the *Rphylopars* package (v 0.2.12) (Goolsby *et al.* 2017), using Pagel’s lambda model of trait evolution. We did this separately for each of the 100 phylogenetic trees, such that each tree was associated with specific imputed values. This method performs well in predicting missing species’ data (Penone *et al.* 2014) and imputed data increase the statistical power of analysis (Nakagawa & Freckleton 2008). Importantly, the bias in imputed data sets tends to be lower than the bias in data sets with missing data omitted, particularly when values for many species are missing (Penone *et al.* 2014). To minimize concerns that imputed data may affect our conclusions, we validated the robustness of our findings by performing all analyses also on the subset of species for which we have data on body mass and sexual size dimorphism.

#### Model specification

We specified two types of models. First, we ran a set of univariate models with peak song frequency as the dependent variable and with either body mass (species or male), sexual size dimorphism (based on wing length or body mass) or habitat density (tree cover or habitat type) as predictor. Second, we ran multivariate models, which included different sets of predictors. The first models included combinations of species body mass, wing-based sexual size dimorphism and tree cover (or habitat type), the second models included combinations of male body mass and body mass-based sexual size dimorphism as predictors. Note that the results from univariate and multivariate models, from analyses based on imputed or raw data, from analyses with species-or male-specific body mass, as well as from analyses based on tree cover or habitat type were qualitatively almost identical (Fig. S2 and Table S1). Hence, in the main text we report only findings from multivariate model containing species-specific body mass, wing-based sexual size dimorphism and tree cover with imputed missing data for body mass and sexual size dimorphism.

## RESULTS

Species-specific median peak song frequency ranged from 215 Hz to 10,659 Hz (*n* = 5,085 species), but most passerine species emitted songs of intermediate frequencies (mean ± SD = 4,030 ± 1,626 Hz; median = 3,790 Hz; Fig. 1a). Median peak song frequency shows a strong evolutionary signal with a coefficient λ ≈ 0.87 (see also Table S1). Nevertheless, low and high peak song frequencies occur within phylogenetically distinct groups (Fig. 1a).

**Figure 1.**
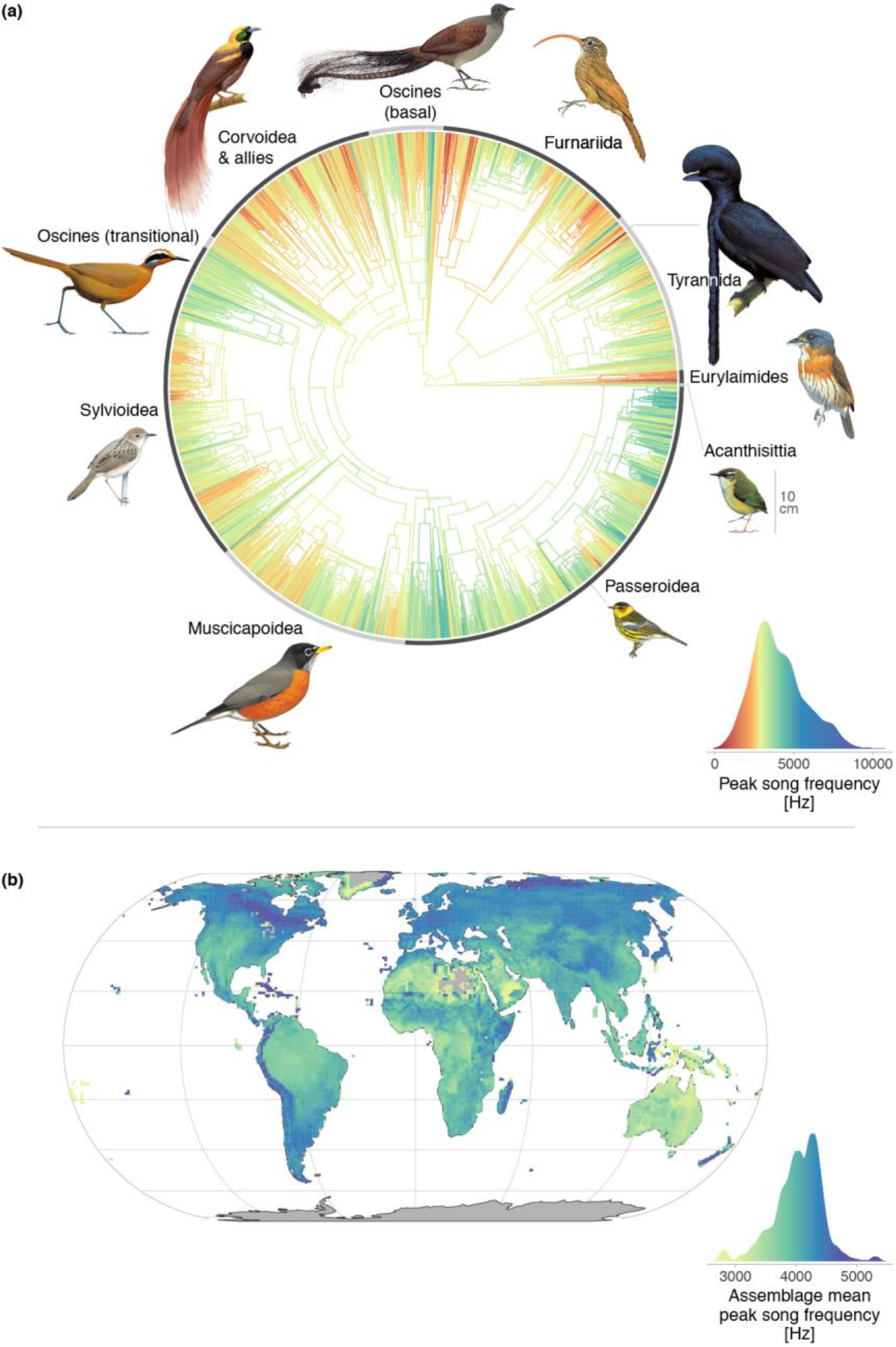
Distribution of peak song frequency across passerines. (a) Distribution across a maximum credibility phylogenetic tree (based on 100 trees sampled from http://birdtree.org) with colour scale reflecting variation (Kernel densities) in species median values (*n* = 5,085 species). Highlighted are 10 major groups of passerines with their representative species, scaled according to size, except for the downscaled representatives of the Tyrannida (should be ~20% larger) and the basal Oscines (should be three times larger); starting with Acanthisittia and going counterclockwise, the pictures depict *Xenicus gilviventris* (10 cm body size), *Smithornis sharpei* (17 cm), *Cephalopterus penduliger* (41 cm; example of low-frequency singer: https://www.xeno-canto.org/75792), *Campylorhamphus trochilirostris* (25 cm), *Menura novaehollandiae* (103 cm), *Paradisaea raggiana* (34 cm), *Eupetes macrocerus* (29 cm), *Cisticola chiniana* (14 cm), *Turdus migratorius* (25 cm) and *Setophaga tigrina* (13 cm; example of high-frequency singer: https://www.xeno-canto.org/182791). Illustrations reproduced by permission of Lynx Edicions. (b) Geographical distribution in peak song frequency across species assemblages (based on the species’ breeding range) defined for 112.5 × 112.5 km (~1° scale) areas. Colour scale reflects variation (Kernel densities) in assembly mean peak song frequency (*n* = 10,856 points; for clearer illustration of differences, outliers were assigned a single value causing the “bumps” on both ends of the distribution).

Passerines sang at low frequencies predominantly in large parts of Australia, in tropical rainforests of the Neotropical, Afrotropical, and Papua New Guinea regions, and possibly in the Sahara where data coverage was sparse (Fig. 1b). Conversely, high-frequency songs characterize passerine communities in the northern parts of the Nearctic and Palearctic regions, in large mountain ranges such as the Andes and Himalayas, in southern parts of the Neotropical region, and in belts of grassland and savannah in Africa (Fig. 1b).

Body mass was the strongest predictor of global variation in peak song frequency (Fig. 2a and Fig. S2), explaining 11-16% of the variation (59-67% together with phylogeny; Table S1). As predicted from the morphological constraint hypothesis, heavier species sang at lower frequencies (Fig. 2a and Fig. S2); this pattern was observed for all but two families (*n* = 52 families with more than 15 species; Fig. 2b and Fig. S3).

**Figure 2.**
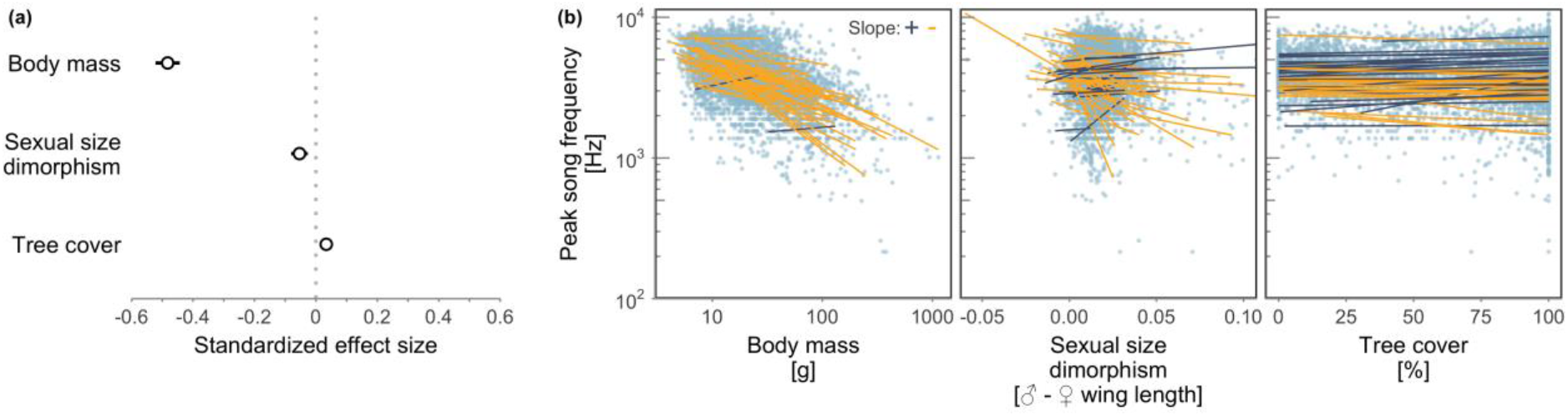
Associations between peak song frequency and body mass, sexual size dimorphism (in wing length) and tree cover across passerines (*n* = 5,085 species). (a) Standardized effect sizes (dots) with their 95% confidence intervals (horizontal lines) based on a multivariate analysis with imputed missing data for body mass and sexual size dimorphism (see Material and Methods and Table S1 for details). Values represent averages from 100 multivariate models, each using a different phylogenetic tree. (b) Relationship between peak song frequency and each of the three explanatory variables. Each dot represents the median peak song frequency of a given species. Lines show the results of univariate robust linear regressions for each of the 52 families with more than 15 species. Positive slopes are indicated in dark blue, negative slopes in yellow. Note the log-scale for peak song frequency and body mass and that for clearer visualisation two lower and ten higher sexual size dimorphism points are not displayed. Robust regressions were fitted to the data with imputed missing values using the rlm function from the *MASS* package (Venables & Ripley 2002). For results of univariate models and those using the original, non-imputed data only, see Fig. S2 and S3, and Table S1.

Peak song frequency was also significantly associated with sexual size dimorphism (either measured in wing length or in body mass), although the effect size was substantially smaller, explaining 1-3% of the variation (Fig. 2a and Fig. S2; Table S1). As predicted based on the sexual selection hypothesis, species with a stronger male-biased sexual size dimorphism (i.e. a higher intensity of sexual selection) sang with lower frequencies, even after controlling for body mass *per se* (Fig. 2a and Fig. S2; Table S1). This effect of decreasing frequency with increasing dimorphism was seen in 67% of families (35 out of 52 families with more than 15 species) while in the remaining families the trend was in the opposite direction (Fig. 2b and S3). Note that in this analysis data on body mass were not sex-specific. Hence, adding sexual size dimorphism might improve model fit, simply because our measure of body mass and sexual size dimorphism together better reflect male size than species-specific mass alone. However, sexual dimorphism in body mass remained influential even when limiting the analysis to a subset of 984 species for which data on male body mass were available (Fig. S2).

Peak song frequency of passerines was weakly, but significantly associated with tree cover or habitat type (Fig. 2a and Fig. S2; Table S1); however, the effect explained only around 0.2% of the variation and was opposite to that predicted from the acoustic adaptation hypothesis: species living in open habitats had lower (not higher) peak song frequencies than those living in more dense, forested habitats (Fig. 2a and Fig. S2; Table S1). Moreover, this effect was observed in only 24 out of 52 families (46%) with more than 15 species (with the random expectation being 50% of the families; Fig. 2b and S3). This unexpected relationship was close to zero and not statistically significant in multivariate models that used the original, nonimputed values of body mass and sexual size dimorphism (based either on wing length or body mass; Fig. S2; Table S1).

## DISCUSSION

Our data revealed remarkable variation in peak song frequency among the world’s passerine birds. Our analyses show that most of the interspecific diversity in peak song frequency can be explained by evolutionary history and by body mass, with an additional effect of sexual size dimorphism as a proxy of the intensity of sexual selection. In contrast, our study does not support the acoustic adaptation hypothesis. Opposite to the prediction, we found at best a weakly positive association between habitat density and peak song frequency. Our results thus indicate that the evolution of peak song frequency in passerines is primarily controlled by morphological constraints, as expected from basic physical principles. We further show that peak song frequency may be shaped by sexual selection, but not by habitat-driven selection to maximize song transmission.

We found that after controlling for phylogeny 11-16% of interspecific variation in peak song frequency of passerines is explained by variation in body mass (Table S1). However, phylogeny also explains some of the variation in body mass (Fig. S1) and in a simple linear regression body mass explains ~27% of the variance in peak song frequency. Together, body mass and phylogeny explained almost 70% of the variation in peak song frequency (Table S1). Our results confirm that body size (estimated as body mass in our study) imposes a strong morphological limit on the production of vocalizations of certain frequencies, presumably through a strong correlation with the length of the vocal tract and the size of the labia in the syrinx (Podos 2001; Suthers & Zollinger 2008; Rodríguez *et al.* 2015). The morphological constraint hypothesis can thus be seen as a kind of “null model” (also see Pearse *et al.* 2018) and it is the remaining variation in peak song frequency that needs explanation.

After accounting for body mass, peak song frequency was lower in species where males were larger than females, i.e. in species with – presumably – stronger sexual selection on males. This result is robust to different ways of analysis (Table S1) and supports the hypothesis that sexual selection has shaped the evolution of song frequency (Greig *et al.* 2013; Hall *et al.* 2013; Geberzahn & Aubin 2014; Linhart & Fuchs 2015; Pearse *et al.* 2018). Our comparative study provides evidence that sexual selection led to low-frequency song performance in many families of passerines, presumably in those where song frequency is indicative of the competitive ability of individuals during male–male interactions (Christie *et al.* 2004; Seddon *et al.* 2004; Price *et al.* 2006). Notably, the songs that departed the most in peak frequency from the expected association with body mass – those of three related species from the Cotingidae family (the Amazonian umbrellabird *Cephalopterus ornatus,* the long-wattled umbrellabird *C*. *penduliger*, and the red-ruffed fruitcrow *Pyroderus scutatus*) – were also those that had the lowest peak frequencies documented for any passerine in our data set (< 260 Hz); their peak frequencies are so low that they partly overlap with the fundamental speech frequencies of humans (100-300 Hz), who are, however, more than 100 times heavier (Baken 1987). The umbrellabirds and their close relatives show high male-biased sexual size dimorphism (compared to other passerines) and a lekking mating system where males display together on traditional “exploded” leks and presumably do not provide parental care (del Hoyo *et al.* 2018). In species that produce substantially lower-frequency songs than predicted from the negative frequency-size relationship, sexual selection may have led to the development of a specific vocal apparatus to produce these sounds (Riede *et al.* 2016), such as the unique pendulous oesophageal vocal sacs that are used as a resonator in umbrellabirds (Sick 1954, see also Riede *et al.* 2015 for a non-passerine example). Although selection for low-frequency sounds may in some cases cause a corresponding change in body size (Fitch 1999), it seems more likely that natural (Woodward *et al.* 2005; Ricklefs 2010) and sexual (Björklund 1990) selection on body size underlies most evolutionary shifts in the song frequency of passerines, with an additional effect of sexual selection on the vocal apparatus.

Despite the theoretical basis and some empirical evidence for a negative association between song frequency and habitat density (Morton 1975; Badyaev & Leaf 1997; Buskirk 1997; Bertelli & Tubaro 2002; Blumstein & Turner 2005; Boncoraglio & Saino 2007), our comparative study provides clear evidence against the acoustic adaptation hypothesis. Peak song frequency across the world’s passerines was, if anything, weakly positively instead of negatively correlated with habitat density. Thus, forest-inhabiting species produced sounds that were higher or similar in peak frequency than those of species living in open areas. While other unmeasured biotic and abiotic characteristics of the environment, including consistent background noise produced by wind, rain, insects or other birds, may drive the evolution of peak song frequencies (reviewed in Brumm & Zollinger 2013), we provide solid evidence that habitat density – as used and widely evaluated in bioacoustic studies – had at best a negligible effect on peak song frequency of passerines. Of course, this does not exclude singing-associated behavioural adaptations of birds that improve signal transmission, such as microhabitat selection during perch-singing or display flights (Menezes & Santos 2020). It is noteworthy that at the intraspecific level, birds can adjust their song frequency to local conditions, but these shifts are relatively minor compared to the interspecific variation in frequency we documented in this study (Slabbekoorn & Peet 2003; Slabbekoorn & den Boer-Visser 2006; Nemeth & Brumm 2010; Brumm & Zollinger 2013).

In conclusion, using data of most passerine species and half of the global avian diversity, our study provides three insights into the evolution of acoustic signals. (1) A strong allometric relationship between body size and peak song frequency imposes a clear limit on the evolution of song frequency. (2) Sexual selection seems to cause departures from this allometric relationship, leading to lower-frequency signals than predicted by body size. Further research into the mechanism (e.g. selection on the structure of the vocal apparatus) is of interest. (3) There is no evidence that species in more dense, forested habitats produce songs of lower frequencies. Our study thus challenges the idea that habitat-dependent selection to maximize sound propagation influences the evolution of signal frequency in songbirds. Future work should focus on the link between song frequency, behaviour during vocal performance (e.g. aerial displays), and habitat properties that influence sound transmission and degradation. In general, our study calls for large-scale empirical studies on acoustic signal frequency in other animal groups as independent replication studies.

## Acknowledgements

We are grateful to all contributors and the administrators of the xeno-canto database (https://www.xeno-canto.org/) and Macaulay Library (https://www.macaulaylibrary.org/). Without their contribution, this study would not have been possible. We thank Barbora Blažková for help with the collection of song recordings of endangered species and Liam Revell for help with visualising phylogenies. This study was supported by the Czech Science Foundation project 17-24782S to TA and by the Max Planck Society to BK.

